# PspA-mediated aggregation protects *Streptococcus pneumoniae* against desiccation on fomites

**DOI:** 10.1101/2023.09.27.559802

**Authors:** Jessica R. Lane, Muralidhar Tata, Rahena Yasmin, Hansol Im, David E. Briles, Carlos J. Orihuela

## Abstract

*Streptococcus pneumoniae* (*Spn*) resides in the nasopharynx where it can disseminate to cause disease. One key *Spn* virulence factor is pneumococcal surface protein A (PspA), which promotes survival by blocking the antimicrobial peptide lactoferricin. PspA has also been shown to mediate attachment to dying epithelial cells in the lower airway due to its binding of cell surface-bound mammalian (m)GAPDH. Importantly, the role of PspA during colonization is not well understood. Wildtype *Spn* was present in nasal lavage elutes collected from asymptomatically colonized mice at levels ∼10-fold higher that its isogenic PspA-deficient mutant (Δ*pspA*). Wildtype *Spn* also formed aggregates in mucosal secretions composed of sloughed epithelial cells and hundreds of pneumococci, whereas Δ*pspA* did not. *Spn* within the center of these aggregates better survived prolonged desiccation on fomites than individual pneumococci and were capable of infecting naïve mice, indicating PspA-mediated aggregation conferred a survival/transmission advantage. Incubation of *Spn* in saline containing mGAPDH also enhanced tolerance to desiccation, but only for wildtype *Spn*. mGAPDH was sufficient to cause low-level aggregation of wildtype *Spn* but not Δ*pspA*. In strain WU2, the subdomain of PspA responsible for binding GAPDH (aa230-281) is ensconced within the lactoferrin (LF)-binding domain (aa167-288). We observed that LF inhibited GAPDH-mediated aggregation and desiccation tolerance. Using surface plasmon resonance, we determined that *Spn* forms multimeric complexes of PspA-GAPDH-LF on its surface and that LF dislodges GAPDH. Our findings have important implications regarding pneumococcal colonization/transmission processes and ongoing PspA-focused immunization efforts for this deadly pathogen.

**IMPORTANCE:** *Streptococcus pneumoniae* (*Spn*) is a dangerous human pathogen capable of causing pneumonia and invasive disease. The virulence factor pneumococcal surface protein A (PspA) has been studied for nearly four decades with well-established roles in pneumococcal evasion of C-reactive protein and neutralization of lactoferricin. Herein, we show that mammalian (m)GAPDH in mucosal secretions promotes aggregation of pneumococci in a PspA-dependent fashion, whereas lactoferrin counters this effect. PspA-mediated GAPDH-dependent bacterial aggregation protected *Spn* in nasal lavage elutes and grown *in vitro* from desiccation on fomites. Furthermore, surviving pneumococci within these aggregates retained their ability to colonize naïve hosts after desiccation. We report that *Spn* binds to and forms protein complexes on its surface composed of PspA, mGAPDH, and lactoferrin. Changes in the levels of these proteins therefore most likely have critical implications on *Spn* colonization, survival on fomites, and transmission.

## INTRODUCTION

*Streptococcus pneumoniae* (*Spn*), also known as the pneumococcus, is a Gram-positive pathobiont that causes disease in the very young, the elderly, those who are immunocompromised, and individuals recovering from or experiencing viral infection of the airway (1–4). *Spn* resides in the nasopharynx, from where it can spread to cause otitis media, pneumonia, bacteremia, and meningitis (5). Accordingly, the pneumococcus is a leading cause of middle ear infections, community-acquired pneumonia, and invasive disease (6, 7). *Spn* is transmitted in saliva and nasal secretions and, therefore, requires physical contact of naïve individuals to infected carriers or exposure to contaminated objects, i.e., fomites (8, 9). Along such lines, pneumococci are not considered to be particularly resilient to external stressors and can be killed by cold, heat, and desiccation (10, 11).

One major virulence factor of *Spn* is pneumococcal surface protein A (PspA). PspA is present in all clinical isolates and is a member of the choline-binding protein family (12). Proteome and transcriptome studies have revealed that PspA is one of *Spn*’s most abundant surface proteins and is produced at all stages of infection (13). Since its discovery in 1986 (14), multiple investigators have demonstrated that PspA is required for *Spn* virulence with deletion mutants attenuated in their ability to cause pneumonia or invasive disease (15–17). Moreover, immunization with recombinant PspA blocked nasal carriage and protected mice against lethal challenge by *Spn* when the infecting strain carried the same version of PspA used for immunization (18–20). On this basis, multiple studies have proven PspA to be a leading candidate for future protein-based vaccines against pneumococcal infection (21–24).

PspA is structurally organized into three major domains: the α-helical domain (αHD), proline-rich domain (PRD), and choline-binding domain (CBD) (25, 26) (**Fig. 1A**). Based on the location of the CBD at the carboxy-terminus, which binds non-covalently to phosphorylcholine residues found on the bacterial cell wall and membrane (25, 27), the αHD and PRD are thought to extend outward through the capsule and be exposed beyond the cell surface. The αHD and PRD of PspA are mosaic in structure with considerable variability in their amino acid sequence, resulting in a size range from 67 to 99 kDa across different strains (28–31). This variability stems from the selective pressure imposed by antigen-specific antibodies that target PspA (32). The mosaic structure of PspA also means that its function varies between strains, depending on the presence or absence of specific subdomains within the αHD and PRD.

**FIG 1.**
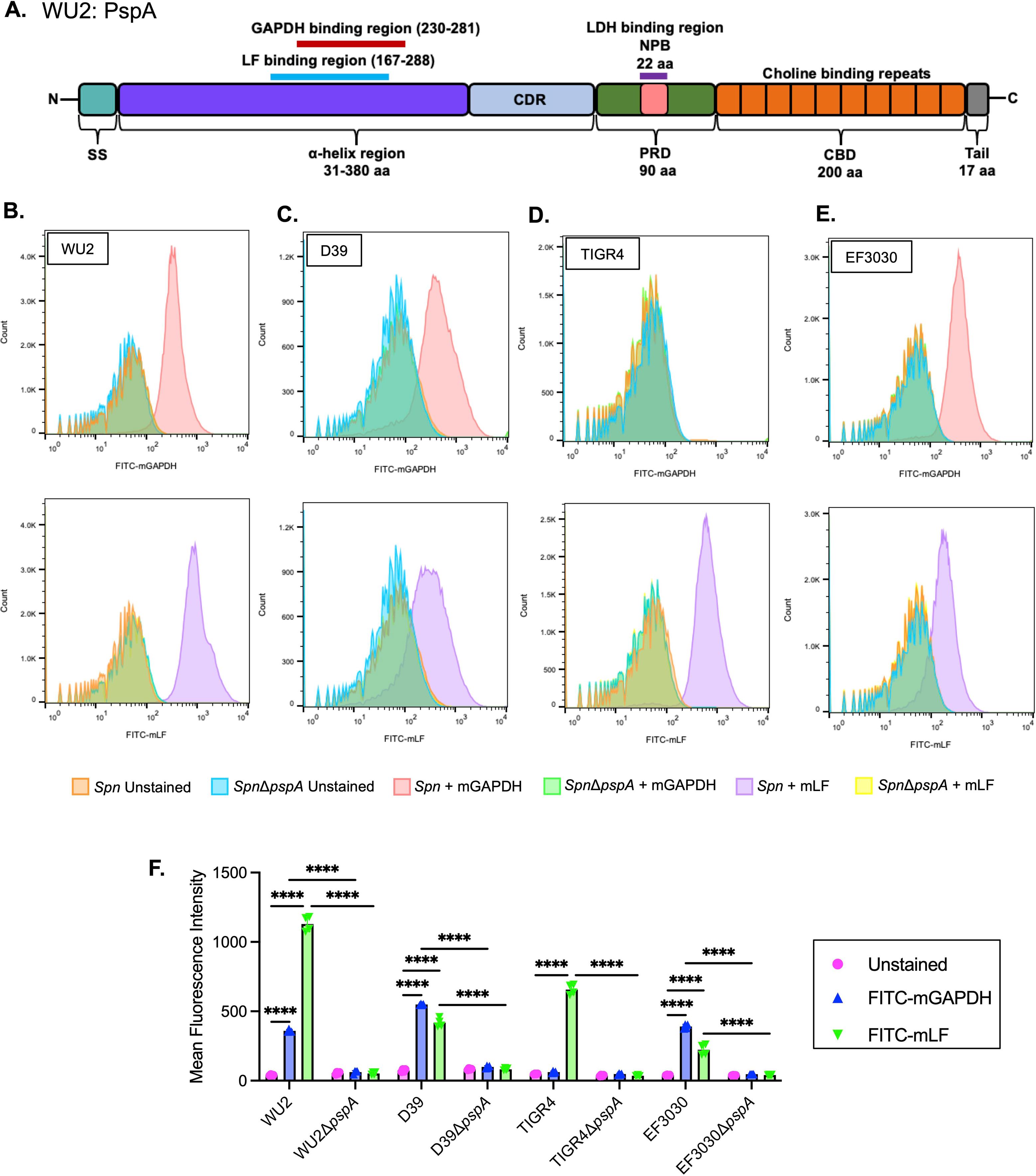
GAPDH and LF bind to *Spn* in a PspA-dependent manner. (A) Structure of WU2 PspA highlighting the α-helix region in purple and the overlapping binding regions of GAPDH (red line) and lactoferrin (LF) (blue line). *Spn* strains (B) WU2 (serotype 3), (C) D39 (serotype 2), (D) TIGR4 (serotype 4), (E) EF3030 (serotype 19F), and their corresponding Δ*pspA* mutants were incubated with FITC-labeled mammalian (m)GAPDH or LF (10 µg/mL) and binding affinity was measured via flow cytometry. (F) Mean fluorescence intensity analysis of FITC-labeled mGAPDH or LF. N=4 with the standard deviation (SD) shown. **** = p ≤ 0.0001.

One major role for PspA is inhibiting recognition by host C-reactive protein (CRP) (33, 34). PspA does this by docking phosphorylcholine residues on the bacterial surface that would otherwise be recognized by CRP (35). This is a general property of all choline-binding proteins, however, PspA has an outsized role in protection due to its abundance. Shaper *et al.* showed that the αHD of PspA also binds to human lactoferrin (LF) and, by means not fully understood, neutralizes the antimicrobial activity of lactoferricin (36). In 2021, our group reported two new roles for PspA. We first described that a conserved 22-amino non-proline block (NPB) found within the PRD domain of some versions of PspA binds to mammalian lactate dehydrogenase A (LDH-A) (37).

Binding resulted in the co-opting of host LDH-A activity, which in turn, enhanced *Spn* virulence when pneumococci were pre-incubated with lactate (37). We also reported that PspA could function as a bacterial adhesin, albeit only to dying host cells. PspA was found to have a specific affinity for mammalian (m)GAPDH, which in turn was adhered to phosphatidylserine residues flipped outward on the cell membrane of lung epithelial cells undergoing programmed cell death (38, 39). The adhesin trait of PspA was shown to be promoted by *Spn*’s pore-forming toxin pneumolysin and was also a feature of secondary pneumococcal infections following Influenza (40). We proposed that this close association mediated by PspA to dying host cells was beneficial to *Spn* as it conferred accessibility to released nutrients that otherwise would be diluted within mucosal secretions, one example being lactate.

Critically, and although the role of PspA during pneumonia and invasive disease is well characterized (14, 16, 34–36, 41), its role during asymptomatic colonization of the nasopharynx remains unclear. Herein, we investigated how PspA-mediated binding to mGAPDH influenced colonization, its shedding, and survival beyond its release from the host. Our results identify a new role for this remarkably versatile protein and reiterate why PspA is vital for pneumococcal persistence in the host.

## RESULTS

For these studies we used serotype 3, strain WU2 as our prototype of *Spn* as well as its version of PspA. Studies with serotype 3 strains are warranted since rates of serotype 3 associated disease have been steadily climbing since 2010 despite the inclusion of the capsule type in current versions of the conjugate vaccine (42, 43). For the sake of comprehensiveness, we also include in our analyses the contribution of PspA in strains belonging to serotype 2, 4, and 19F.

*Spn* binds to both GAPDH and LF in a PspA-dependent fashion.

The αHD of PspA has been reported to bind mGAPDH and LF (37, 38). For *Spn* serotype 3 strain WU2, the mGAPDH binding region of PspA (aa 230-281 of the αHD) is located within the LF-binding region (aa 167-288) (**Fig. 1A**) (41). Using flow cytometry and FITC-labeled proteins, we first validated the ability of wildtype WU2 to bind to recombinant mGAPDH (FITC-mGAPDH) and recombinant LF (FITC-LF). Strong binding to both proteins was observed for wildtype WU2. In contrast, its isogenic *pspA* deficient mutant, i.e., Δ*pspA*, failed to bind mGAPDH or LF (**Fig. 1B and F**). We also tested mGAPDH and LF binding to serotype 2 strain D39, serotype 4 strain TIGR4, and serotype 19F strain EF3030 (**Fig. 1C through F**). Robust mGAPDH binding was observed for D39 and EF3030, but not TIGR4. In contrast, strong LF binding was observed for all three strains. Upon further exploration, we discovered a short in-frame insertion of DNA within the GAPDH-binding motif of TIGR4 PspA that was absent in the other three strains (**Fig. S1**) and propose this is the reason for lack of GAPDH binding. Thus, we confirmed our prior report that WU2 binds to GAPDH (38). But also determined that not all *Spn* strains have this ability, one such exception being the often-used laboratory strain TIGR4.

### PspA-dependent aggregation enhances nasal colonization

Nasal lavage fluid from colonized mice have been shown to contain *Spn* in aggregates, in some instances composed of hundreds of pneumococci, with the latter often bound to host cell debris (44). These aggregates are thought to be biofilm pneumococci formerly or still attached to sloughed mucosal epithelial cells that have been expelled (45, 46). Since *Spn* adheres to dying lung epithelial cells via PspA-mediated interactions (38), we sought to determine if PspA-mediated interactions with mGAPDH were responsible for forming these nasopharynx-derived *Spn*/host cell aggregates. Notably, two days post-inoculation, median bacterial titers in nasal lavage fluid from asymptomatically WU2-colonized mice were ∼10-fold higher than those from Δ*pspA*-colonized mice despite similar starting inoculums (**Fig. 2A**). Microscopic examination of nasal lavage elutes from WU2-colonized mice showed clear and pronounced interactions between pneumococci and sloughed host components, with aggregates comprised of pockets of both live and dead bacteria (**Fig. 2B**). While there were fewer pneumococci in nasal lavage from mice colonized with Δ*pspA*, those observed were individual and showed little to no signs of bacterial aggregation. Subsequently and using ImageJ, we quantified the size of all pneumococci in captured tile scans each 6 X 6 mm of stained nasal lavage elutes. This analysis confirmed that that wildtype-inoculated mice formed significantly larger bacterial aggregates of bacteria compared to *ΔpspA*-inoculated mice (**Fig. 2C**). However, it also showed that *Spn* within nasal elutes were a mixed population and that the majority of CFU were small clusters of bacteria less than 20 µm^2^ in size or individual diplococci (∼2 µm^2^) (**Fig. 2D**). Importantly, we detected mGAPDH via immunoblot in nasal lavage fluid from *Spn* colonized mice as well as from uncolonized mice (**Fig. 2E**). Therefore, while mGAPDH release is not dependent on colonization, its presence in mucosal secretions supports the notion that PspA-mediated adhesion to host cells via mGAPDH was occurring for WU2.

**FIG 2.**
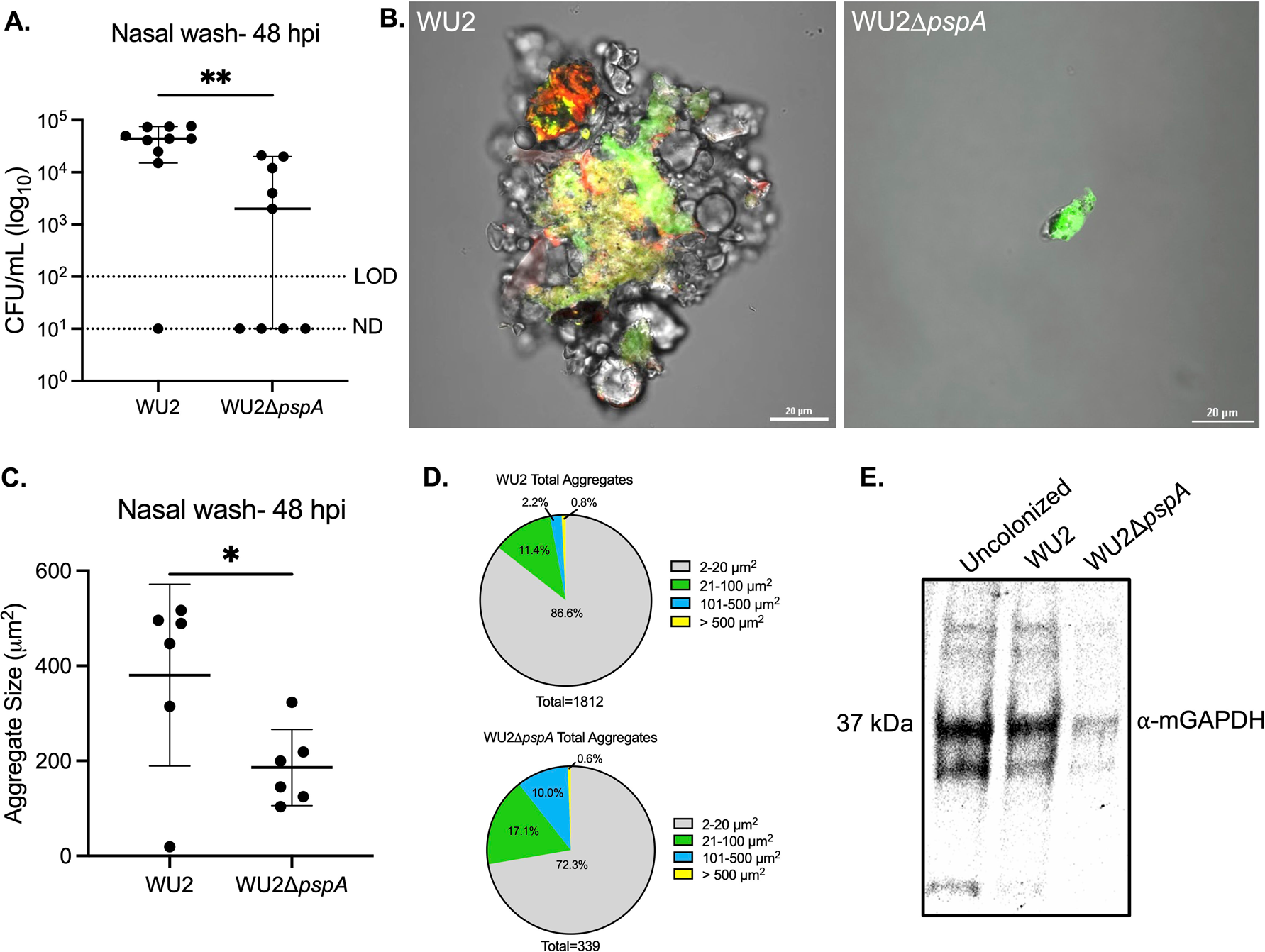
*Spn* aggregates form *in vivo* in a PspA-dependent manner enhancing colonization. 9-week-old C57BL/6J female mice were intranasally inoculated with ∼10 CFU/mL of WU2 (∼6.90 x 10 CFU/mL) or WU2Δ*pspA* (∼7.60 x 10 CFU/mL). (A) Bacterial burden was determined 48 hours post-inoculation (hpi) by colony forming units (CFUs) obtained from nasal washes with saline. N=10 per group and the median with 95% confidence interval is shown. Limit of detection (LOD)=10 CFU/mL. Not detected (ND)= 10 CFU/mL. (B) WU2 aggregate stained with SYTO 9 (green) and propidium iodide (red) collected from nasal wash at 60X magnification under oil immersion and aggregate from mouse inoculated with WU2Δ*pspA* (scale bar=20 μm). (C) Quantification of aggregate size from nasal wash based on live bacteria from the ten largest aggregates (see methods for more detail). N=6 per group with the standard deviation (SD) shown. (D) Total aggregates quantified from nasal washes based on size distribution. (E) Immunoblot of combined nasal wash from uncolonized (n=4), WU2-inoculated (n=5), and WU2Δ*pspA*-inoculated (n=5) mice probed with mouse ⍺-GAPDH (1:200) and IgG-HRP (1:10000). * = p ≤ 0.0332; ** = p ≤ 0.002.

### PspA protects *Spn* against desiccation ex vivo

Due to greater access to nutrients and, presumably, encasement within a protective matrice, we hypothesized that PspA-mediated aggregation with sloughed nasopharyngeal cells would confer to *Spn* prolonged tolerance to desiccation following expulsion in mucosal secretions. Nasal lavage elutes from WU2 and Δ*pspA*-colonized mice were spotted onto glass slides, air-dried at room temperature; then, after 24-, 48-, and 72-hours, submerged in saline for 30 seconds, gently scraped off the glass, and plated onto agar plates to determine if recoverable bacteria were present and, if so, enumerate the percent survival (**Fig. 3**). Starting at 24 hours, we observed a strong trend for a greater rate and number of recoverable bacteria by WU2 versus Δ*pspA*. This became significant at 48 hours but then declined by 72 hours when only ∼20% of the bacteria were recoverable for either cohort. Subsequently, we sought to determine if *Spn* that did survive desiccation could colonize a naïve host. To answer this question, we used nasal elute recovered from colonized mice that had been desiccated for 48 hours to inoculate the nares of naïve mice after rehydration in saline. Half the mice inoculated with formerly desiccated WU2 had recoverable *Spn* in the nasopharynx 48 hours after inoculation. In contrast, all mice inoculated with desiccated nasal elutes from WU2Δ*pspA*-colonized mice remained uncolonized (**Fig. 4**). Together these observations indicate PspA is involved during colonization/transmission processes and prolongs bacterial infectivity *ex vivo* via attachment to sloughed host cells, aggregate formation, and resistance to desiccation-mediated killing.

**FIG 3.**
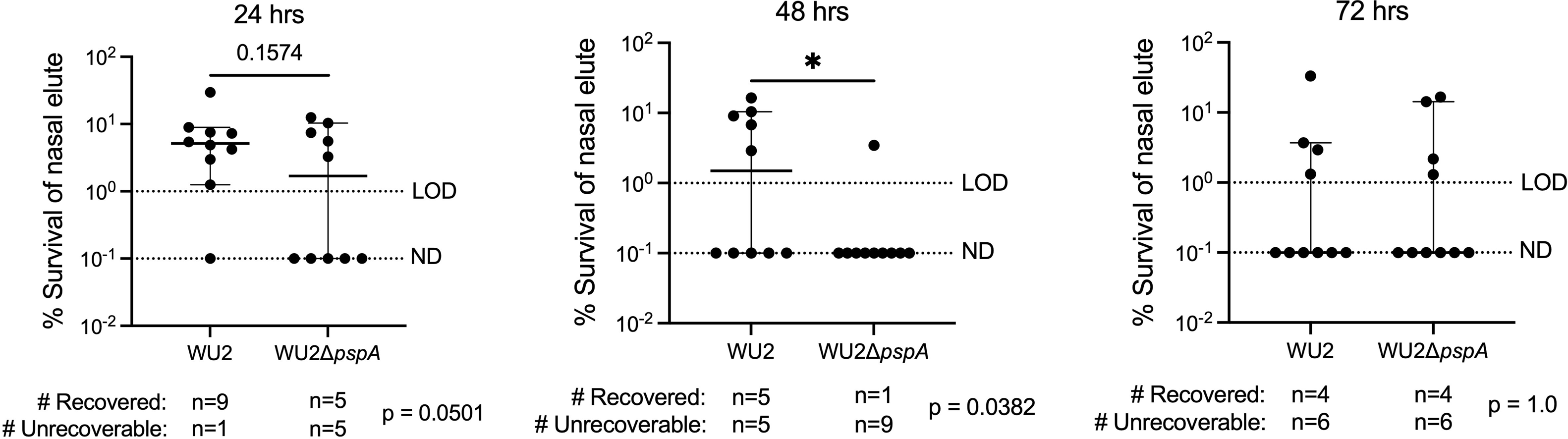
PspA confers a protective capability to *Spn* against desiccation ex vivo. 9-week-old C57BL/6J female mice were intranasally inoculated with ∼10 CFU/mL of WU2 (∼7.40 x 10 CFU/mL) or WU2Δ*pspA* (∼8.30 x 10 CFU/mL), as previously described. Nasal wash elutes were plated onto glass slides and allowed to dry out. Bacteria were rehydrated and survival was enumerated after 24-, 48-, and 72-hours. The number of recoverable bacteria per treatment after 24-, 48-, and 72-hours is shown with the Chi p-value calculated below the graphs. Starting bacterial burden from nasal lavage of WU2 (∼4.50 x 10 CFU/mL) and WU2Δ*pspA* (∼3.80 x 10 CFU/mL) colonized mice. N=10 per group and the median with a 95% confidence interval is shown. LOD=10 % survival. ND=10 % survival. * = p 0.0332.

**FIG 4.**
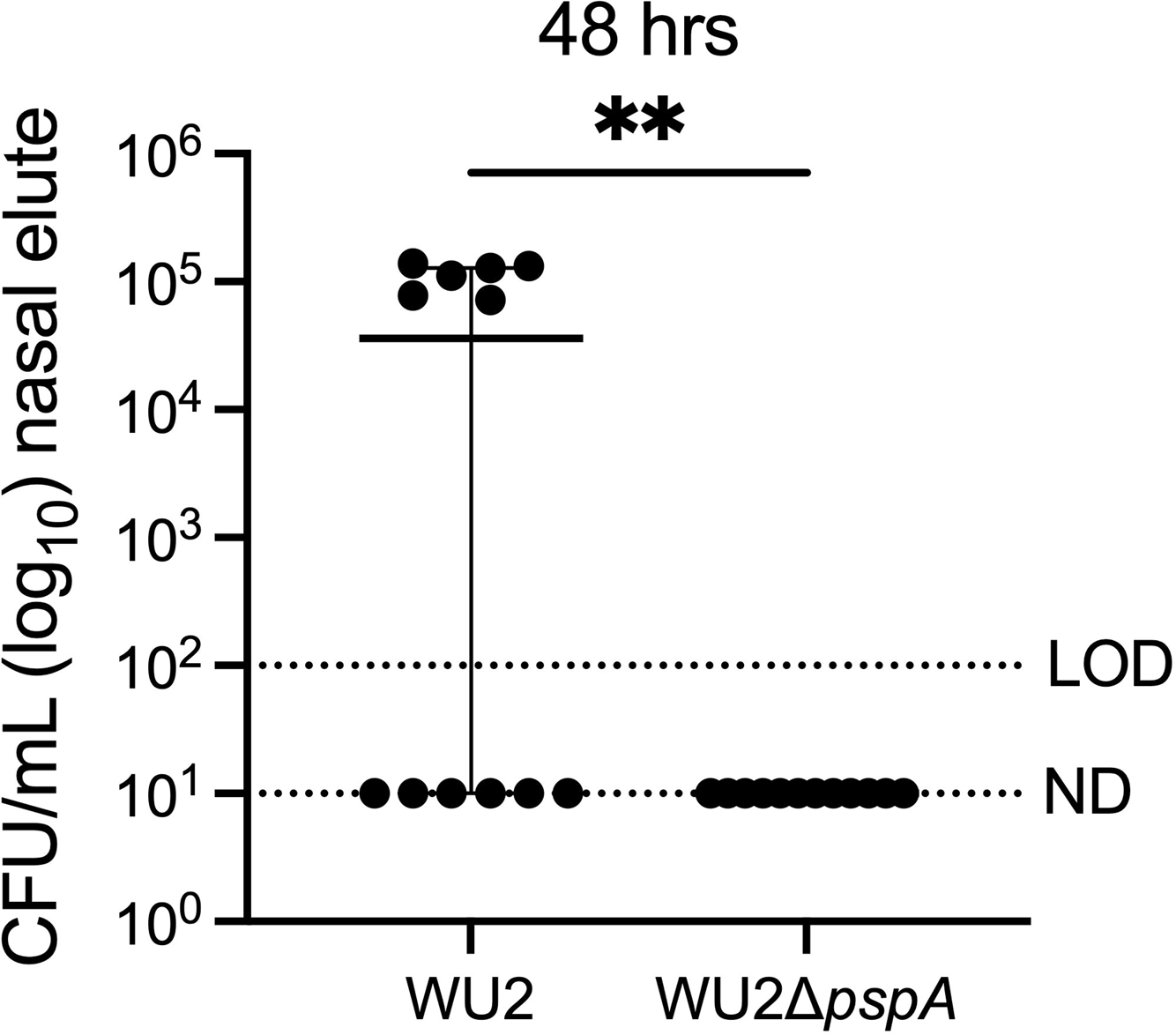
*Spn* remains infectious after desiccation in a PspA-dependent manner. 9-week-old C57BL/6J female mice were intranasally inoculated with ∼10 CFU/mL of WU2 (∼3 x 10 CFU/mL) or WU2Δ*pspA* (∼4.2 x 10 CFU/mL), as previously described. After 48 hours, nasal wash elutes were plated onto glass slides and allowed to dry out. After 48 hours of desiccation, nasal elutes were rehydrated and inoculated intranasally into naïve 6-week-old C57BL/6J female mice. Bacterial burden was determined 48 hpi by CFUs obtained from nasal washes with saline. N=12 per group and the median with a 95% confidence interval is shown. LOD=10^2^ CFU/mL. ND= 10^1^ CFU/mL. ** = p ≤ 0.002.

### GAPDH-binding confers protection against desiccation in vitro

Given these results, we sought to determine whether GAPDH was sufficient to confer PspA-mediated desiccation tolerance or if instead other host-factors were required. To do this, we incubated WU2 and Δ*pspA* grown to exponential phase in THY media with 10 μg/ml mGAPDH and, following our desiccation protocol, re-examined survival after 24-, 48-, and 72-hours. Across all time points, we observed that WU2 incubation with mGAPDH was sufficient to confer protection against desiccation compared to untreated bacteria (**Fig. 5A; Table S1**). This trait was confirmed as being PspA-dependent as Δ*pspA* had significantly lower survival at any time points tested. Live/dead staining of rehydrated WU2 showed that most pneumococci were dead following desiccation (**Fig. S2**), however, when survivors were present they were within clusters of pneumococci that formed at a frequency of <1% of CFU. Notably, incubation of WU2 with LF did not enhance survival against desiccation. When WU2 was added to saline containing mGAPDH and LF there was also no protection; the latter suggesting LF impeded the protection afforded by GAPDH. Consistent with this result, D39 mixed with GAPDH had a significant survival advantage versus D39Δ*pspA* after 7 and 14 days and EF3030 was trending towards greater survival than EF3030Δ*pspA* after 14 days. TIGR4, which does not bind GAPDH, did not have increased survival against desiccation and were all killed after 3 days (**Fig. S3**). Thus, we observed a very wide range in *Spn* resistance to desiccation across multiple strains which was enhanced by the ability to bind GAPDH.

**FIG 5.**
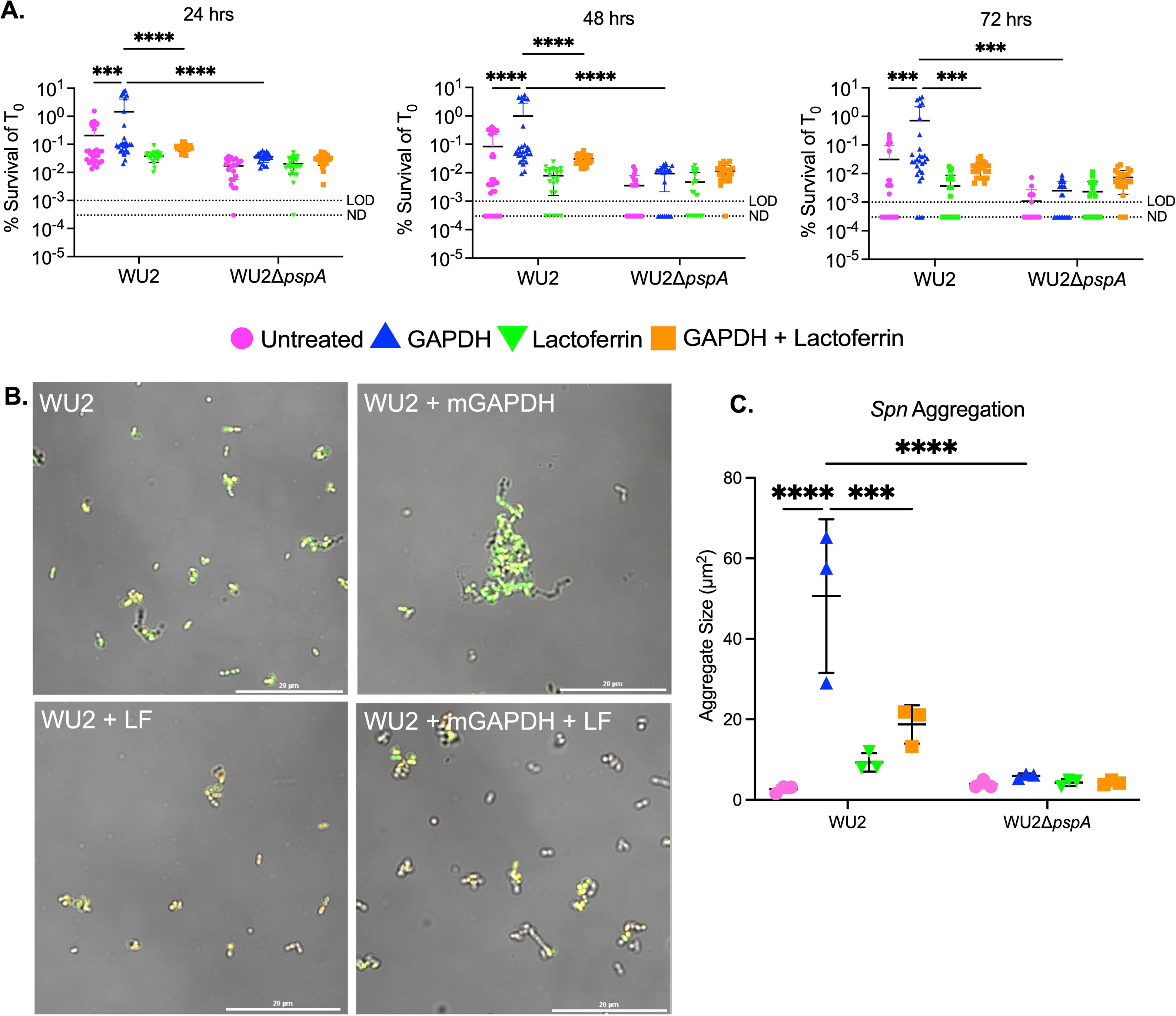
*Spn* forms PspA-dependent aggregates in vitro that protect against desiccation. (A) WU2 and WU2Δ*pspA* were incubated with mGAPDH, LF, or with mGAPDH and LF (10 µg/mL) and plated onto glass slides and allowed to dry for 24-, 48-, and 72-hours. Bacterial survival was enumerated by CFUs based on T_0_ survival (∼3.0 x 10 CFU/mL). N=24-29 with the standard deviation (SD) shown. LOD=10 % survival. ND=3 x 10 % survival. (B) High resolution image of WU2 pneumococci stained with SYTO 9 (green) and propidium iodide (red) and fixed with 4% paraformaldehyde and Fluoromount™ (see methods for more details). WU2 shown when incubated in solution with mGAPDH, LF, or with mGAPDH and LF (10 µg/mL). All images captured at 60X magnification under oil immersion with a 20 μm scale bar (see methods for more details). (C) Mean cluster size of the five largest clusters per treatment (10 µg/mL) quantified from live bacteria (see methods for more details). N=3 with the standard deviation (SD) shown. *** = p ≤ 0.0002; **** = p ≤ 0.0001.

### *Spn* is aggregated by GAPDH in the presence of PspA

Our observation that mGAPDH conferred desiccation resistance by itself raised the question of whether it was doing so by aggregation. We tested this by enumerating the maximal size of clusters that formed following treatment of WU2 with different concentrations of mGAPDH and LF (**Fig. S4A**). Importantly, we saw that treatment of WU2 with 10 μg/mL mGAPDH for 30 minutes caused the formation of pneumococcal clusters at a low but consistent frequency of <1% of CFU (**Fig. 5B and C; Fig. S4B**). Interestingly, this did not occur at lower or higher concentrations. No cluster formation was observed with the Δ*pspA* mutant mixed with mGAPDH (**Fig. S4C**). Similarly, we tested the impact of LF and saw no clustering. What is more, mixture of *Spn* with 10 μg/mL of mGAPDH and LF blocked the formation of these clusters (**Fig. 5C**), an observation supporting the importance of aggregation on desiccation survival and the antagonistic role of LF on this feature. Aggregation with mGAPDH was also confirmed with D39 and EF3030, albeit at lower cluster sizes (**Fig. S5**). Unexpectedly, the same result was observed for TIGR4, suggesting that GAPDH may bind to other factors on the bacterial surface. This is evident by the aggregation of TIGR4Δ*pspA* at levels slightly higher than wildtype. D39Δ*pspA* and EF3030Δ*pspA* may also contain other GAPDH-binding factors as they too bound GAPDH at levels slightly lower or equal to their wildtype counterparts, respectively. Finally, adding either GAPDH or LF to WU2 or WU2Δ*pspA* in nutrient media had no impact on growth (**Fig. S6**).

### GAPDH and LF bind concurrently to PspA and form a multimeric complex

Realizing the importance of the PspA-GAPDH interaction *in vivo* and *ex vivo*, we wanted to more directly test how LF affected *Spn*’s interactions with WU2 PspA by seeing if GAPDH and LF bound simultaneously to the bacterial surface or if one protein was favored. WU2 exposed simultaneously to FITC-mGAPDH and Alex647-labeled LF (Alexa 647-LF) were all double positive for these markers following flow cytometry (**Fig. 6A**). Pre-incubation of WU2 with LF showed no impact on the subsequent binding of mGAPDH, while pre-incubation of WU2 with mGAPDH had a nominal effect on LF binding (**Fig. 6B**). These results suggest that pneumococci can become decorated with both host proteins simultaneously, with no significant differences or interference in the capability to do so. To further interrogate the interactions between these proteins, we examined the ability of recombinant WU2 (r)PspA possessing only αHD and PRD to bind mGAPDH, LF, or both using surface plasmon resonance. When mGAPDH was injected over rPspA-amine coupled to a sensor chip followed by LF, or vice versa, we observed sequential binding of both proteins onto rPspA (**Fig. 6C**). The sensograms also demonstrated that the complex’s dissociation slope differed depending on which protein was bound to PspA first (**Fig. 6D**). These differences could be attributed to the known dissociation constant (Kd) of the interacting partners: PspA-mGAPDH is 2.95 µM (38); PspA-LF is 3.89 ±2.74 nM (**Fig. 6E**); mGAPDH-LF is 43.8 ±8.26 nM (41). As the Kd of PspA-LF is stronger than mGAPDH-LF, it is therefore expected that mGADPH, when followed by LF, would dissociate faster from PspA in comparison to LF followed by mGAPDH (**Fig. 6D inset**). Thus, LF could ultimately displace GAPDH from PspA preventing the aggregation of *Spn* and the protection it affords.

**FIG 6.**
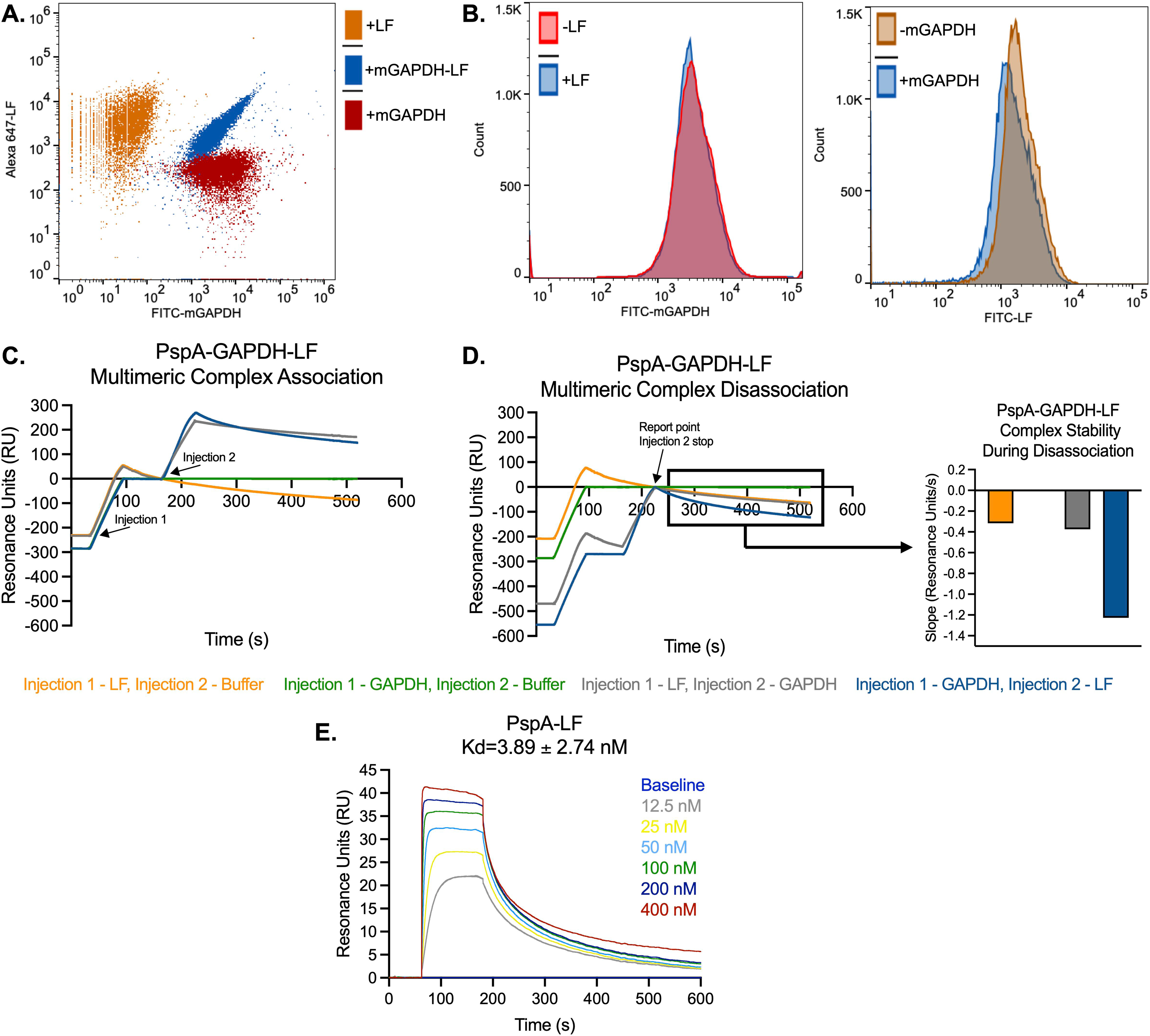
PspA-GAPDH-LF form a multimeric complex. (A) WU2 was co-incubated with FITC-labeled mGAPDH and Alexa 647-labeled LF and binding affinity for both proteins to PspA was measured via flow cytometry. (B) The two histograms represent the individual binding affinities for mGAPDH and LF with and without the addition of the other protein. (C) Surface plasmon resonance (SPR) plot of the association and complex formation of GAPDH and LF to rPspA amine-coupled to a sensor chip. (D) SPR plot of the disassociation of the PspA-GAPDH-LF complex. Inset image is the stability constant during disassociation. (E) The disassociation rate constant (Kd) of PspA and LF.

## DISCUSSION

*Spn* has been studied for over a century with several virulence factors that help it evade the host immune system identified and characterized during that time. PspA is one such factor with established and critical roles of blocking opsonization by C-reactive protein, preventing lactoferricin-mediated killing, adhesion to dying host cells, and co-opting of host factors for its metabolic benefit having been previously described (33, 34, 36–38). Multiple and diverse roles for surface proteins are not uncommon amongst bacterial pathogens and most likely reflect evolutionary pressure toward optimal utilization of these proteins. Notably, strain-specific properties for PspA have previously been described. For example, we reported that *Spn* binds to lactate dehydrogenase only when PspA has a non-proline block present within its PRD (37). Herein, we demonstrate that the TIGR4 version of PspA does not bind GAPDH, most likely due to an insertion in its α-helical domain that was absent in the other strains tested. This variability in surface protein function is not restricted to PspA as some versions of choline-binding protein A (CbpA) bind to Factor H, whereas others do not (47). Likewise, pneumococcal serine-rich repeat protein (PsrP), which binds to cytokeratin 10 and to itself, varies dramatically in predicted size from 37 to >2000 kDa (48, 49). This diversity in function/structure reflects the mosaic nature of these proteins with distinct subdomains having specific properties.

Our previous studies showed that PspA and host GAPDH interactions during pneumococcal infection impacted bacterial localization in the lower airway; specifically, that PspA allowed pneumococci to bind dying epithelial cells in the lungs (38). However, the importance of this interaction in the upper respiratory tract (URT), i.e. the nasopharynx, remained unexplored. Our group has also previously reported that pneumococci in the nasopharynx form aggregates/biofilms on mucosal epithelial cells that can contain hundreds of bacteria and are sloughed during nasal lavage (44). This study not only reaffirmed this observation, but also showed that PspA plays an essential role in their formation. It is notable that other pneumococcal surface proteins also promote aggregation. For example, deletion of PsrP in serotype 4 strain TIGR4 abrogated aggregate formation in nasal lavage fluid from colonized mice (44). Notably, WU2 and EF3030 carry low amounts of PsrP while D39 does not carry any (49), suggesting that for these strains PspA may have a compensatory function. This was observed in our aggregation and desiccation experiments with these strains and GAPDH. Munoz-Elias *et al.* screened a transposon library for *Spn* mutants unable to form biofilms on polystyrene and found that 23 of 29 mutants identified as biofilm-deficient were subsequently impaired for nasopharyngeal colonization (50). Virulence genes identified by Munoz-Elias as being vital included the pneumococcal pilus, choline-binding protein F, and CbpA (50). Thus, promoting aggregation/biofilm formation *in vitro* seems to be a convergent property of pneumococcal surface proteins and promotes colonization.

Desiccation is an ever-present threat to bacteria residing in the nasopharynx. Known mechanisms for bacterial tolerance to desiccation stress include replacing water hydrogen bonds with trehalose, scavenging reactive oxygen species, and supporting bacterial architecture with an extracellular polysaccharide matrix (51–54). In this study, we showed that PspA-mediated bacterial aggregation provided moderate resistance to desiccation. Prior work by our group has shown that capsular polysaccharide can act as an antioxidant and confers resistance to oxidative stress by *Spn* (55). PspA-mediated GAPDH-dependent aggregation of pneumococci may shield those in the center of the aggregate from oxidative stress that occurs during colonization and potentially provide them with nutrients released by attached dying host cells. Such a scenario is consistent with our detection of viable pneumococci within aggregates following live/dead staining.

Marks *et al.* showed that pneumococci within *in vitro*-grown biofilms better survive desiccation than their planktonic counterparts (11). Additionally, survival of pneumococci within biofilms has been recorded for up to 20 days on a dry, inanimate surface (56, 57). Our results suggest the length which pneumococci can survive desiccation, ranging from less than 2 days up to 2 weeks, is highly strain dependent. However, when PspA binds GAPDH, there was a notable improvement. Further exploration is required using isogenic capsule switch variants, as well as isogenic PspA switch variants, to dissect the contribution of capsule, PspA, and other encoded factors on bacterial resistance to desiccation.

We initially presumed that PspA-mediated aggregation was exclusively the result of its binding to immobilized mGAPDH bound to the surface of dying host cells with the dying host cell acting as a surface to which multiple bacteria could bind. Instead, we observed that mGAPDH also mediates infrequent, but measurable, levels of bacterial aggregation on its own and that this low level of aggregation is sufficient to promote survival of *Spn* following desiccation. Thus, *Spn* carrying versions of PspA that bind GAPDH most likely benefit from both; the latter enabling the bacteria to bind to each other within nasal secretions. Along such lines, it is important to consider that LF blocked aggregation *in vitro* and, therefore, possibly serves as an inhibitor to pneumococcal aggregation *in vivo*. LF itself is a multi-functional molecule with potent antimicrobial properties, is primarily produced by neutrophils, and its presence in mucosal secretions increases as inflammation intensifies (58, 59). LF production by the host therefore also serves to disrupt pneumococcal aggregation and, in turn, enhance bacterial susceptibility. Future studies are warranted to determine how the interplay between *Spn-*mediated mucosal epithelial cell damage, mGAPDH release, and levels of LF influence aggregation and the duration of *Spn* colonization.

As part of our study, we sought to explore the consequence of the overlapping binding regions of PspA’s mGAPDH- and LF-binding motifs. We expected that these molecules would compete with one another for exclusive binding to PspA. However, we were instead surprised by the observation that PspA bound to both mGAPDH and LF simultaneously. Our results, which showed sequential binding of mGAPDH to PspA-LF and, alternatively, LF to PspA-mGAPDH, as well as work by Rawat *et al.* that show mGAPDH and LF bind to one another (60, 61), together suggest that a multimeric protein complex is formed. Notably, mGAPDH is known to from homodimers (62). Therefore, the scenario where PspA (from bacteria 1)::mGAPDH1::mGAPDH2::PspA (from bacteria 2) is a potential explanation for how mGAPDH is able to mediate aggregation. As LF blocks aggregation, we propose that LF binding to mGAPDH or PspA obscures the ability of the PspA homodimers to interact with PspA molecules on another bacterium and eventually promotes their disassociation. The stoichiometry and structure of this complex is an area of interest to us. Future efforts to resolve this will include creating recombinant mGAPDH and LF that do not bind to one another nor form heterodimers.

In summary, we have identified a new feature that promotes pneumococcal survival following desiccation – PspA-mediated GAPDH-dependent aggregation. This finding has meaningful implications regarding the known roles of PspA during colonization and possibly transmission via survival on fomites. Importantly, its effects can be countered by host LF. Considerable work is needed to fully characterize this new role, including the specific amino acid sequence that is responsible, identification of the domains by which mGAPDH and LF bind to PspA, the structure of this multi-protein complex, the specific facets of aggregation that provide tolerance to desiccation, and the consequence of antibody against PspA in mucosal secretions. We recognize that the phenotypes presented here may differ slightly *in vitro* compared to *in vivo*, and many factors come into play once in a living organism. However, our data presented here adds to the growing literature suggesting that PspA plays a vital role in pneumococcal pathogenesis and a potential role in the transmission of *Spn*.

## MATERIALS AND METHODS

### Ethics Statement

Animal experiments were performed in compliance with the University of Alabama at Birmingham Institutional Animal Use and Care Committee (IACUC). The protocol used in this study (#22157) was approved by the IACUC at the University of Alabama at Birmingham.

### Protein purification

Purification of recombinant proteins PspA (rPspA), GAPDH (mammalian-GAPDH and *Spn*-GAPDH) was performed by using cobalt-affinity resin and the *E. coli strain* (NEBExpress® Iq) or BL21(DE3) carrying the respective plasmids. The strains were grown at 37°C in Luria-Bertani (LB) broth with required antibiotics and expression was induced at 0.4 OD_621_ by the addition of 1 mM IPTG. After 4 hours of induction, cells were harvested and lysed in BugBuster Master Mix (MilliporeSigma, 71456-4) with 1 mM PMSF for 30 minutes at room temperature with gentle shaking. The cell lysates were spun at 14,000 RPM for 20 minutes and the supernatant was loaded on beads pre-equilibrated with default buffer (5 mM Imidazole, 50 mM Tris-HCl pH 7.5, 150 mM NaCl). Beads were washed 10 times with default buffer and protein was eluted in default buffer containing 100 mM Imidazole. Collected protein fractions were checked on an SDS-PAGE gel and concentrated/buffer exchanged to PBS using a centrifugal concentrator tube. Protein concentration was measured by using Pierce™ BCA Protein Assay Kit (ThermoFisher, 23225).

### Fluorophore labeling of proteins

FITC and Alexa Fluor 647 labelling of purified proteins was done by using FluoroTag FITC Conjugation Kit (MilliporeSigma, FITC1-1KT) and Alexa Fluor™ 647 Antibody Labeling kit (Invitrogen, A20186), respectively, following the manufacturer’s recommendation.

### Flow cytometry

*Spn* was cultured to an OD_621_ of 0.4 (∼6.0 X 10^7^ CFU/mL) and harvested after centrifugation (3,500 g for 10 minutes). *Spn* was washed twice in FACS buffer (PBS, 3% FBS, and 0.1% sodium azide) and resuspended in the same buffer to approximately 5.0 X 10^6^ CFU/mL. 100 μL of cells were incubated at room temperature with 10 μg/mL of FITC-labeled mGAPDH, Alexa Fluor 647 labeled LF, or both. After 30 minutes, cells were washed and resuspended in FACS buffer and analyzed using flow cytometry with 100,000 events collected (BD, AccuriTHC6 Flow Cytometer). Mean fluorescence intensity was calculated using FlowJo (version 10) from n=4 with both technical and biological replicates. Strains used: WU2, D39, TIGR4, EF3030, and their corresponding isogenic Δ*pspA* mutants (Table S2).

### Surface plasmon resonance (SPR)

The multimeric complex formation of PspA-GAPDH-LF was determined by using surface plasmon resonance (Cytiva, Biacore T200 instrument). PspA (ligand) was amine-coupled to a CM5 sensor chip at a concentration of 3 µg/mL by setting the maximum immobilization to 50 resonance units (RU), following the manufacturer’s instructions. Binding analysis was performed at 25°C by injecting 25 nM of LF followed by mGAPDH or mGAPDH followed by LF, diluted in PBSP+ running buffer (0.02 M phosphate buffer with 2.7 mM KCl, 0.137 M NaCl and 0.05% Tween 20), over PspA and a blank surface. The blank surface was used as a control for monitoring non-specific binding and performing reference subtraction. The surfaces were regenerated between injections with 3 M magnesium chloride. All experiments were done in duplicate. The experimental binding data was analyzed using the Biacore T200 Evaluation Software version 2.0 according to the manufacturer’s instructions. Each experiment was performed 3 times with our figure showing a representative run. For kinetic studies, LF was diluted in PBSP+ at indicated concentrations and injected over PspA immobilized on a CM5 chip as described above. The experimental data was analyzed using the Biacore T200 Evaluation Software 2.0 according to the manufacturer’s instructions. The experimental data for PspA-LF interactions was fitted to a two-state binding curve with a Kd value of 3.89 nM (±2.74).

### Bacterial aggregation

*Spn* was cultured in Todd-Hewitt broth with yeast (THY) and grown to exponential phase before being resuspended in PBS at an OD_621_ of 0.1 (∼3.0 X 10^7^ CFU/mL). 0.5 mL aliquots were incubated with 10 μg/mL or increasing concentrations (0-100 μg/mL) of recombinant human GAPDH or LF for 30 minutes at 37°C. Bacteria were stained with SYTO 9 and propidium iodide specific for prokaryotes (LIVE/DEAD® BacLight™ Bacterial viability Kit, Molecular Probes L7012) according to the manufacturer’s instructions for 15 minutes at room temperature and no light. After a brief vortex, 20 μL was spotted onto a clean glass slide and fixed with 4% paraformaldehyde (PFA) and Fluoromount™ Aqueous Mounting Medium (Sigma, F4680) before a cover slip was added. Representative images shown were captured using a confocal microscope at 60X magnification under oil immersion (scale bar is 20 μm) (Nikon A1R). The size of the aggregates was calculated from images of live aggregated *Spn* captured using a Leica LMD6 with DFC3000G-1.3-megapixel monochrome camera at 40X magnification (Leica Biosystems, Buffalo Grove, IL). The top five largest aggregates were determined using ImageJ (NIH) quantification of live bacteria with three replicates per treatment.

### *In vivo* nasal colonization and desiccation

C57BL/6J female mice aged ∼9 weeks were inoculated intranasally with *Spn* strain WU2 and its isogenic Δ*pspA* mutant (∼1 X 10^6^ CFU/mL). After 48 hours, 10 μL of PBS was used to wash the nasal cavity of each mouse while under isoflurane sedation. ∼2 μL of recovered liquid was then diluted ten-fold, mixed thoroughly with a pipette, and plated onto 5% sheep’s blood agar plates for colony forming unit (CFU) enumeration. Recovered bacteria from the nasal washes were stained as described above. Representative images shown were captured using a confocal microscope at 60X magnification under oil immersion as described above (scale bar is 20 μm) (Nikon A1R). Tile scan images (6×6 mm) were captured for each sample using a Leica LMD6 with DFC3000G-1.3-megapixel monochrome camera at 10X magnification (Leica Biosystems, Buffalo Grove, IL). Size of live bacterial aggregates was based on the top ten largest aggregates obtained using ImageJ (NIH) as described above. As each mouse had multiple aggregates ranging from the very small (<4 μm^2^) to the very large (> 400 μm^2^), we chose the largest aggregates from the nasal elutes for quantification as to focus on the formation and not frequency of the aggregates. The inoculation was repeated and the recovered liquid from the nasal wash was then plated onto glass sides as described for the desiccation experiments below.

Percent survival was calculated in the same manner as described below. For the fomite infectivity experiment, the same protocol as above was followed. After 48 hours of desiccation, nasal elutes were rehydrated and roughly ∼10-15 μL of liquid was inoculated into the nares of naïve ∼6-week-old C57BL/6J female mice. Nasal washes with PBS were done at 24-, 48-, and 72-hours post-inoculation for CFU enumeration.

### Immunoblots of nasal lavage

Recovered nasal lavage fluid from asymptomatically colonized mice was treated with NuPAGE™ LDS Sample Buffer (4X) (ThermoFisher, NP0007) and beta-mercaptoethanol before baking at 95°C for 10 minutes. Samples were loaded onto an SDS-PAGE gel (4-15%) before transferring on to a nitrocellulose membrane. After blocking with 5% BSA, the membrane was incubated overnight with the primary antibody followed by washing with 1% TBS-Tween20 and incubation with the secondary antibody. Visualization of the blots was done using the Pierce™ ECL Western Blotting Substrate (ThermoFisher, PI32209) and ChemiDoc XRS+ System (BioRad). Primary antibody: rabbit α-mouse GAPDH (Santa Cruz: sc-32233) (1:200). Secondary antibody: goat α-mouse IgG-HRP serum (1:10,000).

### Bacterial desiccation

*Spn* was cultured and treated as stated above. After a brief vortex, 50 μL was spotted onto a clean glass slide and air dried for 24-, 48-, and 72-hours at room temperature. Dried samples were resuspended in 50 μL of PBS and dilutions were plated onto 5% sheep’s blood agar plates for CFU enumeration. Survival percentage was calculated by dividing desiccated CFU by the time zero (T_0_) CFU taken right after the initial incubation (∼3.0 x 10^6^ CFU/mL).

### Statistical analysis

All statistical analyses were done using GraphPad Prism (version 9). For comparisons of two groups, the nonparametric one-tail or two-tailed Mann-Whitney *t*-test was used and the median with a 95% confidence interval or the standard deviation (SD) is shown. For all multiple comparisons, a nonparametric 2-way analysis of variance (ANOVA) with Tukey’s pairwise test was used, or a Chi^2^ test when indicated, and the standard deviation (SD) is shown.

## ACKNOWLEDGEMENTS

JRL, MT, and RY designed and performed the experiments. HI helped design and provided feedback on the experiments and results. CJO designed, supervised, and conceptualized the project and experiments. JRL and CJO wrote and edited the manuscript. DEB provided essential feedback on the manuscript. This work was supported by the National Institutes of Health (NIH) grant: 5R01AI156898-03.

