## Supplemental data for "PspA-mediated aggregation protects *Streptococcus pneumoniae* against desiccation on fomites"

**Table S1. Recovery of pneumococci on fomites following desiccation (n=24-29/cohort).**

| 24 Hours | Recovered | Unrecoverable | % Recovered | P-value ($Chi^2$) |
|---|---|---|---|---|
| **Untreated** | | | | |
| WU2 | 29 | 0 | 100 | 0.0056 |
| WU2Δ*pspA* | 16 | 5 | 76 | |
| **GAPDH** | | | | |
| WU2 | 29 | 0 | 100 | NA |
| WU2Δ*pspA* | 21 | 0 | 100 | |
| **Lactoferrin** | | | | |
| WU2 | 24 | 0 | 100 | 0.1486 |
| WU2Δ*pspA* | 22 | 2 | 92 | |
| **G + LF** | | | | |
| WU2 | 24 | 0 | 100 | 0.3122 |
| WU2Δ*pspA* | 23 | 1 | 96 | |

| 48 Hours | Recovered | Unrecoverable | % Recovered | P-value ($Chi^2$) |
|---|---|---|---|---|
| **Untreated** | | | | |
| WU2 | 12 | 16 | 43 | 0.3046 |
| WU2Δ*pspA* | 6 | 15 | 29 | |
| **GAPDH** | | | | |
| WU2 | 28 | 0 | 100 | 0.0004 |
| WU2Δ*pspA* | 13 | 8 | 62 | |
| **Lactoferrin** | | | | |
| WU2 | 14 | 10 | 58 | 0.0417 |
| WU2Δ*pspA* | 7 | 17 | 29 | |
| **G + LF** | | | | |
| WU2 | 24 | 0 | 100 | 0.1486 |
| WU2Δ*pspA* | 22 | 2 | 92 | |

| 72 Hours | Recovered | Unrecoverable | % Recovered | P-value ($Chi^2$) |
|---|---|---|---|---|
| **Untreated** | | | | |
| WU2 | 8 | 20 | 29 | 0.0332 |
| WU2Δ*pspA* | 1 | 20 | 5 | |
| **GAPDH** | | | | |
| WU2 | 25 | 3 | 89 | $\leq$ 0.0001 |
| WU2Δ*pspA* | 5 | 16 | 24 | |
| **Lactoferrin** | | | | |
| WU2 | 7 | 17 | 29 | 0.0645 |
| WU2Δ*pspA* | 2 | 22 | 8 | |
| **G + LF** | | | | |
| WU2 | 22 | 2 | 92 | 0.0330 |
| WU2Δ*pspA* | 16 | 8 | 67 | |

**Table S2. Strains used.**

| Strain | Serotype | Genotype | Reference |
|---|---|---|---|
| WU2 | 3 | Clinical isolate | Hollingshead, Becker et al. 2000 (32) |
| WU2Δ*pspA* | 3 | Isogenic deletion mutant of full length gene of *pspA*, *ermR* | Park et al. 2021 (37) |
| D39 | 2 | Clinical isolate | Lanie et al. 2007 (63) |
| D39Δ*pspA* | 2 | Unmarked in-frame deletion of *pspA* gene by allelic-exchange using the Janus cassette | This study |
| TIGR4 | 4 | Clinical isolate | Tettelin et al. 2001 (60) |
| TIGR4Δ*pspA* | 4 | Unmarked in-frame deletion of *pspA* gene by allelic-exchange using the Janus cassette | This study |
| EF3030 | 19F | Clinical isolate | Mukerji et al. 2012 (35) |
| EF3030Δ*pspA* | 19F | Isogenic deletion mutant of full length gene of *pspA*, *ermR* | Park et al. 2021 (37) |
| MT15 | | NEB®Express Iq Competent *E. coli*/ pQE30-SS01 | Park et al. 2021 (38) |
| MT51 | | BL21-DE3/pET30-2-GAPDH (human) | Park et al. 2021 (38) |

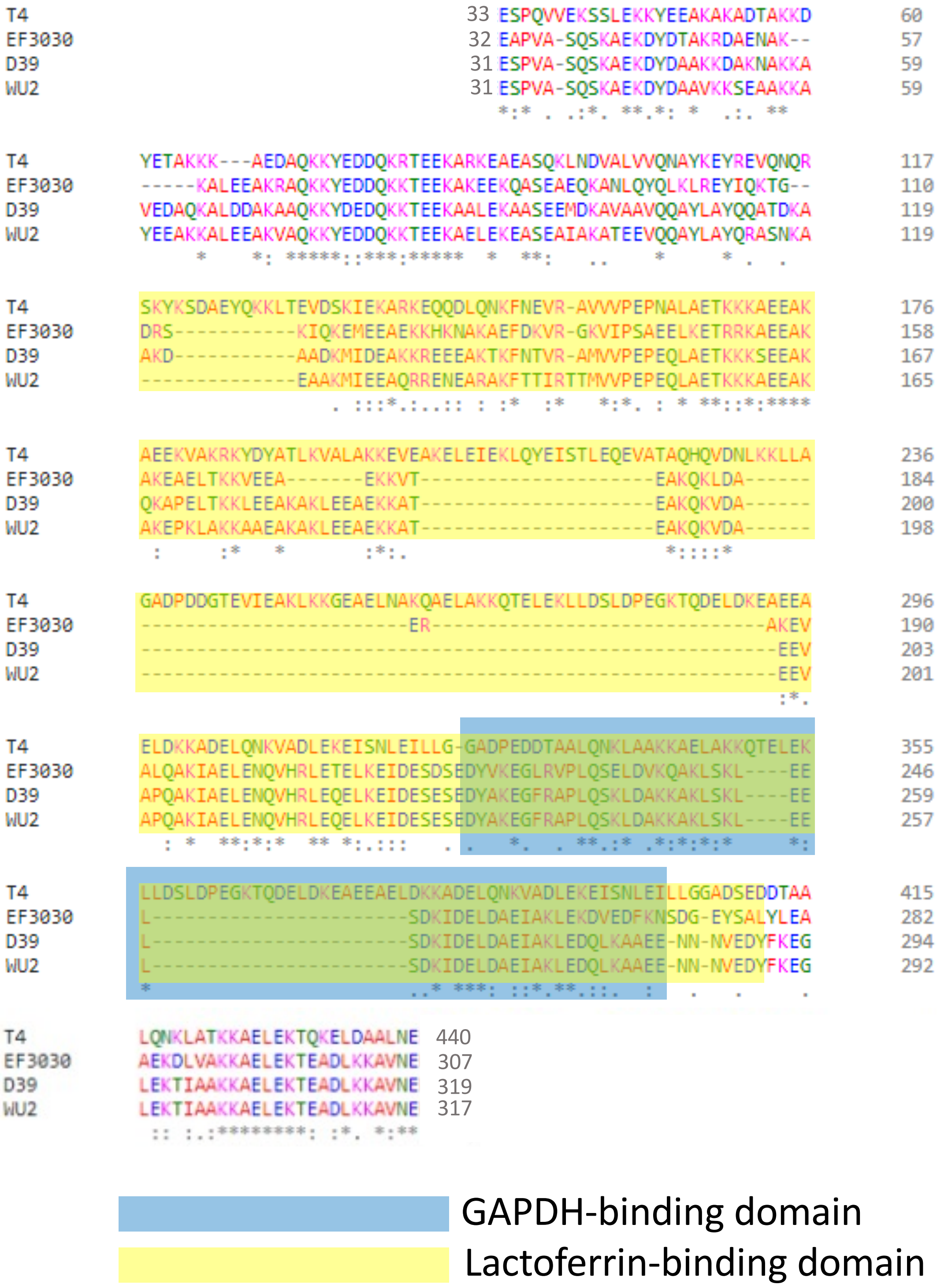


**FIG S1** GAPDH and LF-binding domains of PspA across different *Spn* strains. Multiple sequence alignment of amino acid sequences of PspA for TIGR4 (serotype 4), EF3030 (serotype 19F), D39 (serotype 2), and WU2 (serotype 3). GAPDH-binding domain (blue) and LF-binding domain (yellow) are highlighted within the α-helical region.

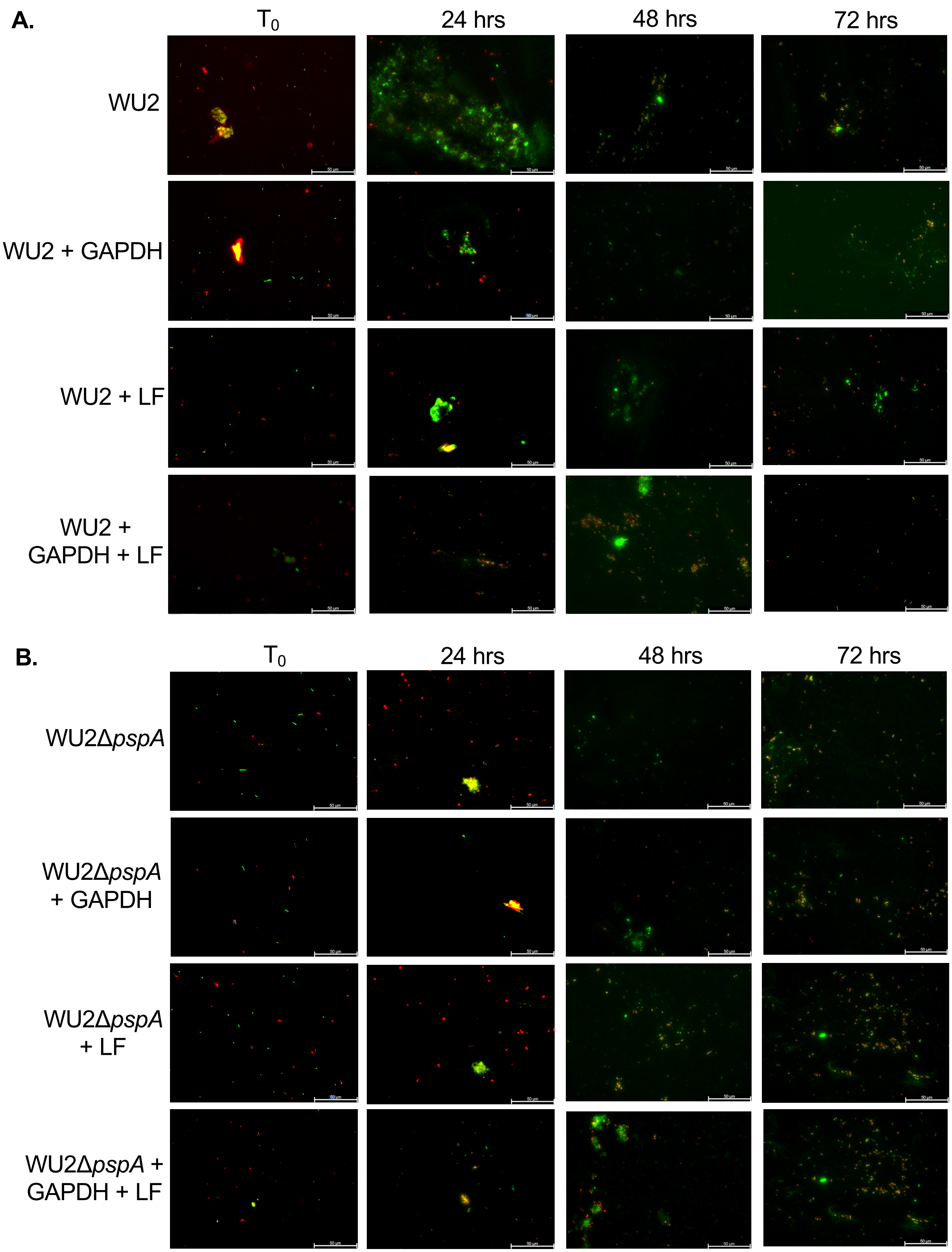


**FIG S2** PspA-GAPDH complex protects *Spn* from desiccation *in vitro* over time. (A) WU2 and (B) WU2Δ*pspA* both in suspension ($T_0$) and desiccated were stained with SYTO 9 (green) and propidium iodide (red). Representative images for time “zero” ($T_0$) aggregated *Spn* and 24-, 48-, and 72-hours post desiccation *Spn* were taken at 10X magnification (scale bar=50 μm).

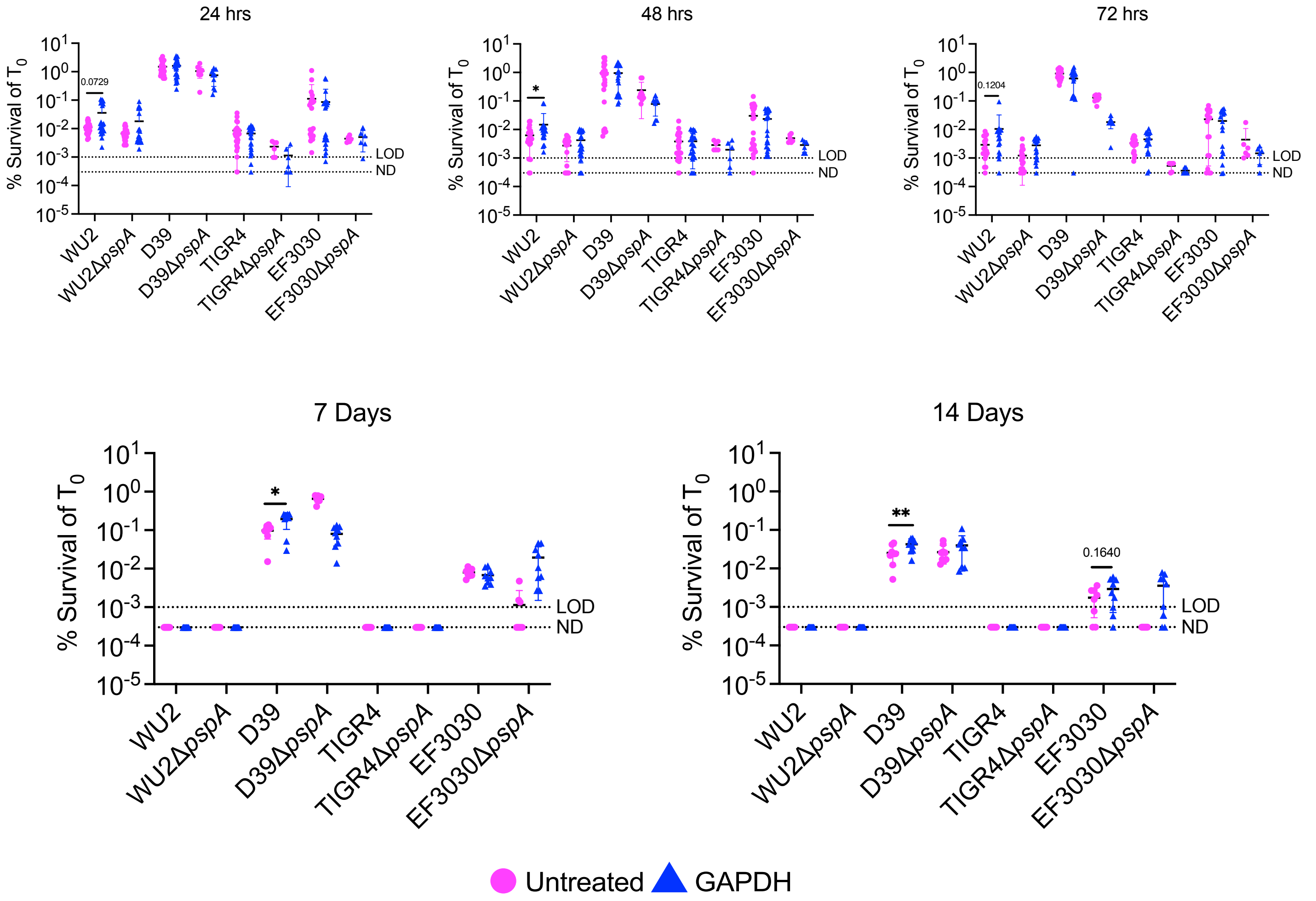


**FIG S3** In vitro desiccation protection over time with multiple *Spn* strains. *Spn* strains WU2 (serotype 3), D39 (serotype 2), TIGR4 (serotype 4), EF3030 (serotype 19F), and their corresponding Δ*pspA* mutants were incubated with mGAPDH (10 µg/mL) and plated onto glass slides and allowed to dry for 24-, 48-, and 72-hours, 7 days, and 14 days. Bacterial survival was enumerated by CFUs based on $T_0$ survival. N=9-24 with the standard deviation (SD) shown. LOD=$10^{-3}$ % survival. ND = 3 x $10^{-4}$ % survival. * = $p \leq 0.0332$; ** = $p \leq 0.002$.

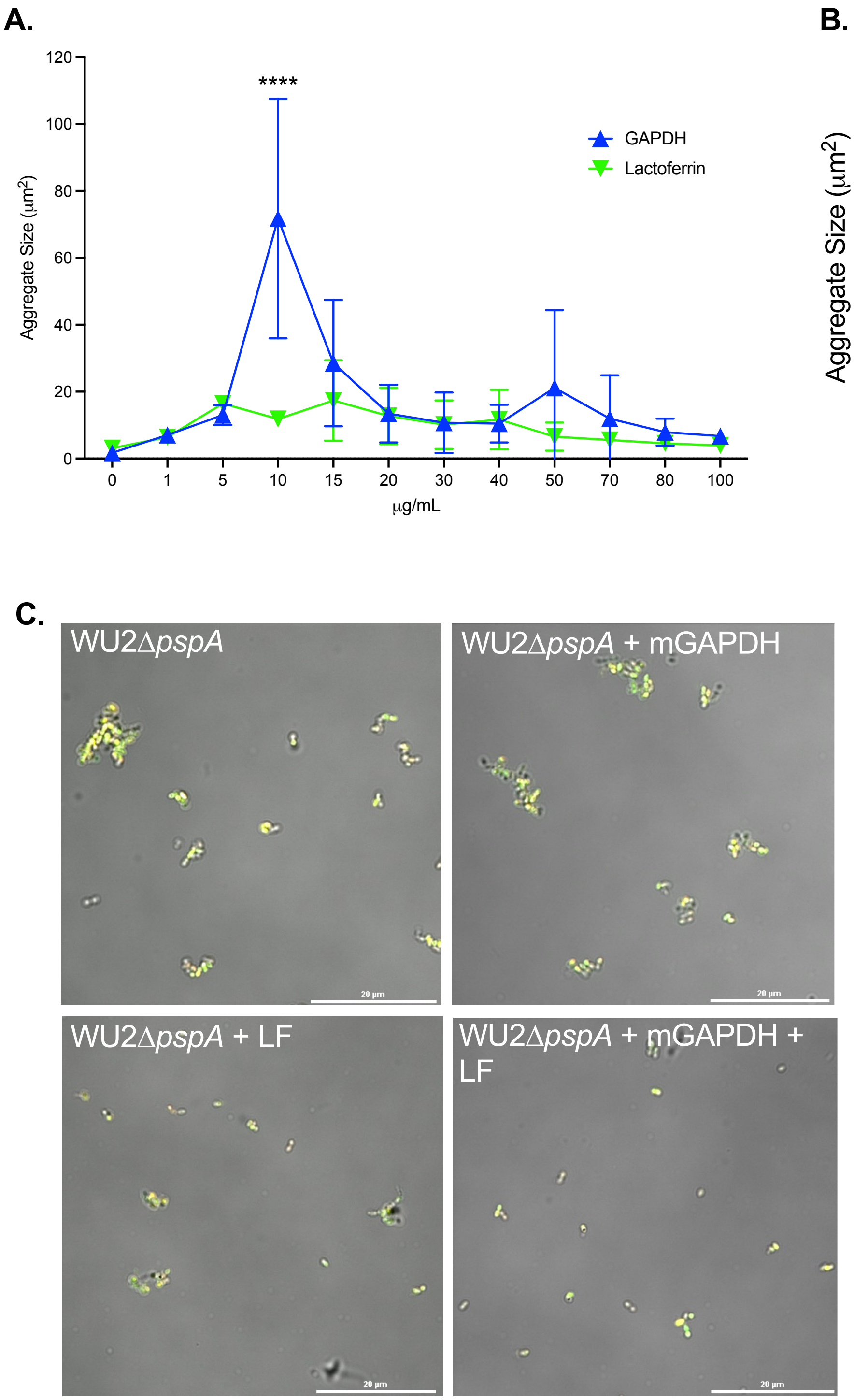


**FIG S4** *Spn* is aggregated by GAPDH in a PspA-dependent manner. (A) Mean cluster size of the five largest clusters of WU2 per treatment with various concentrations of mGAPDH or LF (0-100 µg/mL) quantified from live bacteria. N=3 with the standard deviation (SD) shown (see methods for more details). (B) WU2 was incubated in solution with mGAPDH (10 µg/mL) over time. Mean cluster size of the five largest clusters of WU2 per timepoint was quantified from live bacteria. N=3 with the standard deviation (SD) shown (see methods for more details). (C) High resolution image of WU2Δ*pspA* pneumococci stained with SYTO 9 (green) and propidium iodide (red) and fixed with 4% paraformaldehyde and Fluoromount™ (see methods for more details). WU2Δ*pspA* shown when incubated in solution with mGAPDH, LF, or mGAPDH and LF (10 µg/mL) . All images captured at 60X magnification under oil immersion with a 20 µm scale bar (see methods for more details). * = $p \leq 0.0332$; **** = $p \leq 0.0001$.

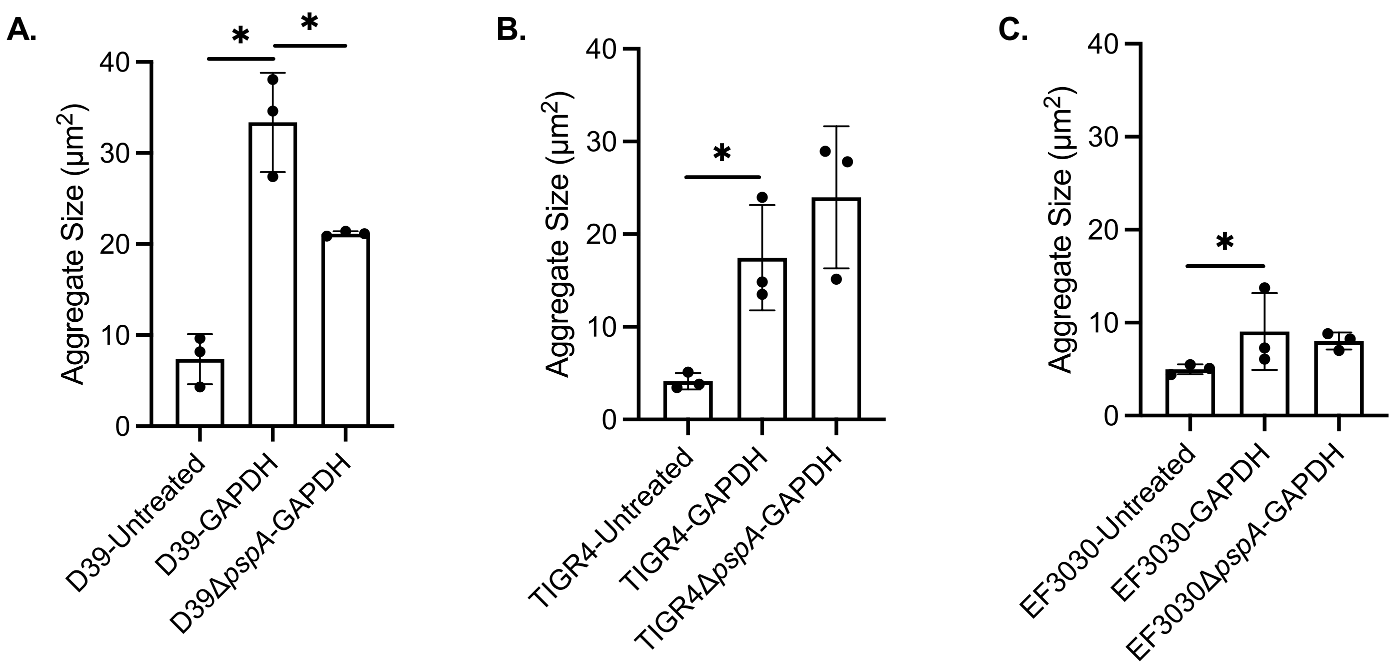


**FIG S5** *Spn* is aggregated by GAPDH in a strain-dependent manner. *Spn* strains (A) D39 (serotype 2), (B) TIGR4 (serotype 4), (C) EF3030 (serotype 19F), and their corresponding Δ*pspA* mutants were stained with SYTO 9 (green) and propidium iodide (red) after incubation with and without mGAPDH (10 μg/mL) (see methods for more details). Mean cluster size of the five largest clusters per treatment (10 μg/mL) were then quantified from live bacteria (see methods for more details). N=3 with the standard deviation (SD) shown. * = p ≤ 0.0332.

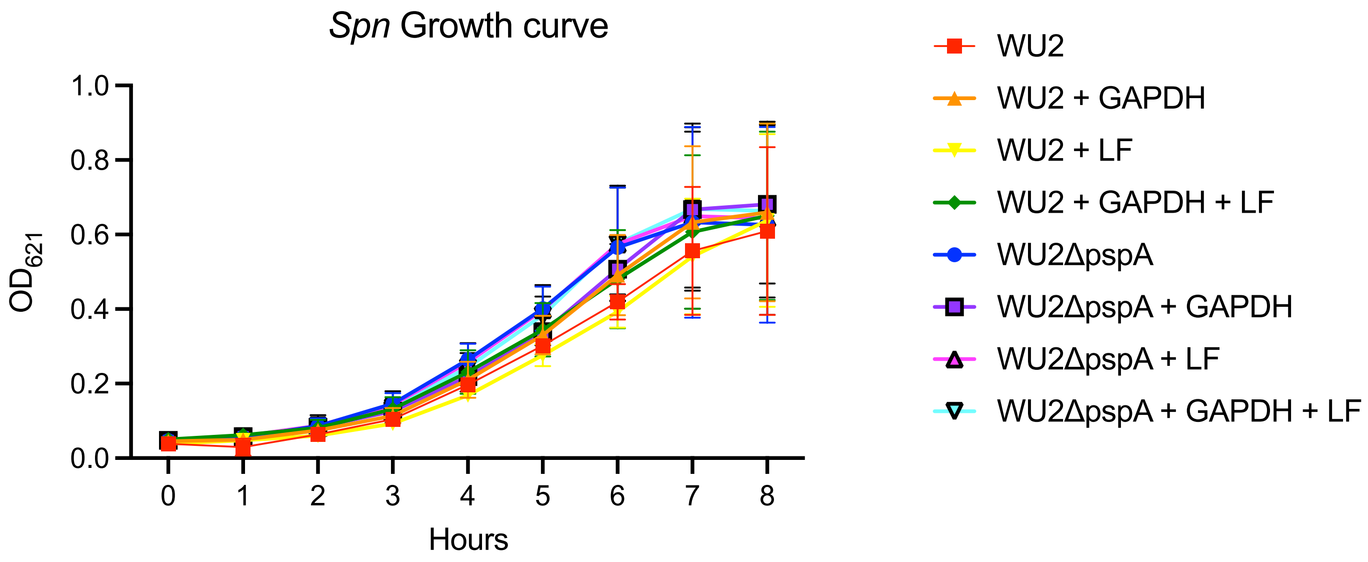


**FIG S6** *Spn* growth is not affected by addition of GAPDH or LF. WU2 and WU2Δ*pspA* were grown in Todd Hewitt-Yeast (THY) broth with or without the addition of mGAPDH, LF, or with mGAPDH and LF (10 μg/mL) in an incubator at 37℃ with 5% $CO_2$ over 8 hours. $OD_{621}$ was measured at each hour to record growth. N=3 with the standard deviation (SD) shown.
